# Polygenic risk scores applied to a single cohort reveal pleiotropy among hundreds of human phenotypes

**DOI:** 10.1101/203257

**Authors:** Adam Socrates, Tom Bond, Ville Karhunen, Juha Auvinen, Cornelius A. Rietveld, Juha Veijola, Marjo-Riitta Jarvelin, Paul F. O’Reilly

**Affiliations:** MRC Social, Genetic and Developmental Psychiatry Centre, Institute of Psychiatry, Psychology and Neuroscience, King’s College London, London, UK; Department of Epidemiology and Biostatistics, MRC-PHE Centre for Environment & Health, School of Public Health, Imperial College London, Norfolk Place, W2 1PG, London, UK; Center for Life Course Health Research, Faculty of Medicine, University of Oulu, Oulu, Finland.; Medical Research Center Oulu, Oulu University Hospital and University of Oulu, Oulu, Finland.; Department of Applied Economics, Erasmus School of Economics, Erasmus University Rotterdam, Rotterdam, 3062 PA, The Netherlands.; Department of Epidemiology, Erasmus Medical Center, Rotterdam, 3015 GE, The Netherlands.; Erasmus University Rotterdam Institute for Behavior and Biology, Rotterdam, 3062 PA, The Netherlands.; Department of Psychiatry, Research Unit of Clinical Neuroscience, University of Oulu, Oulu, Finland; Department of Psychiatry, Oulu University Hospital, Oulu, Finland; Biocenter Oulu, P.O. Box 5000, Aapistie 5A, FI-90014, University of Oulu, Finland; Unit of Primary Care, Oulu University Hospital, Kajaanintie 50, P.O. Box 20, FI-90220 Oulu 90029 OYS, Finland

## Abstract

**Background:** There is now convincing evidence that pleiotropy across the genome contributes to the correlation between human traits and comorbidity of diseases. The recent availability of genome-wide association study (GWAS) results have made the polygenic risk score (PRS) approach a powerful way to perform genetic prediction and identify genetic overlap among phenotypes.

**Methods and findings:** Here we use the PRS method to assess evidence for shared genetic aetiology across hundreds of traits within a single epidemiological study – the Northern Finland Birth Cohort 1966 (NFBC1966). We replicate numerous recent findings, such as a genetic association between Alzheimer’s disease and lipid levels, while the depth of phenotyping in the NFBC1966 highlights a range of novel significant genetic associations between traits.

**Conclusions:** This study illustrates the power in taking a hypothesis-free approach to the study of shared genetic aetiology between human traits and diseases. It also demonstrates the potential of the PRS method to provide important biological insights using only a single well-phenotyped epidemiological study of moderate sample size (~5k), with important advantages over evaluating genetic correlations from GWAS summary statistics only.

## Introduction

The emergence of large-scale GWAS results has demonstrated an enrichment of genetic variants affecting multiple phenotypes, confirming that pleiotropy is a common feature of the human genome [1,2]. Several statistical genetics methods have been developed to quantify this shared genetic architecture formally, such as bivariate genome-wide complex trait analysis (GCTA), LD Score regression and the polygenic risk score (PRS) approach [3–5]. Applying these methods, a large number of studies have tested shared genetic aetiology between two traits, and more recently these have been expanded to estimate pairwise genetic overlap across multiple traits [6–8].

The PRS approach utilises GWAS summary statistics to produce individual-level risk or profile scores and is, therefore, the technique that offers most hope for future personalised or precision medicine [9,10]. Single nucleotide polymorphisms (SNPs) associated with a phenotype below a specific *P*-value threshold are used to produce a score that predicts the risk of that clinical outcome or infers its trait values. Individual polygenic scores can then be used to predict other traits in a regression across a study sample to expose genetic overlap between traits. The key benefits of using the PRS method over alternatives relate to modelling flexibility and statistical power. Exploiting individual-level cohort data allows a greater number of phenotypes and models to be tested [11], relative to relying on GWAS summary statistics only. This can enable greater exposure of causal pathways than via application of LD Score Regression [6]. Moreover, GCTA requires large-scale (*N* > 10k) individual-level data on all tested phenotypes for sufficient power, while LD Score Regression requires large-scale GWAS summary statistics on all phenotypes, yet the PRS method is well-powered from using only large-scale GWAS data for the base phenotypes and relatively small-scale (*N* < 10k) individual-level data on the target traits. A potential limitation of PRS analyses is that subject overlap between the GWAS samples and the target cohort samples can lead to false positive associations. However, this can be addressed directly by recalculating the base GWAS with overlapping cohorts removed, or mathematically by using their GWAS results in a recalculation of the meta-analysis GWAS excluding their effect (see **Methods**).

For the current analysis, PRS were generated in the NFBC1966 participants in relation to 48 phenotypes with large-scale GWAS results available. These PRS were used to predict over 100 traits in the NFBC1966 cohort data. The NFBC1966 contains detailed information on clinical outcomes and health-related behaviour, offering an opportunity to test many traits and combinations not previously investigated. Hence, the depth of phenotyping in the NFBC1966 offer a unique possibility to shed further light on the genetic overlap among phenotypes.

## Methods

5404 participants with genotype data on 364,590 SNPs were available in the NFBC1966 [12]. Baseline data were collected on maternal and offspring demographic, clinical and anthropometric traits in early life. Follow-up data were collected at 14 and 31 years of age on a range of traits, including blood pressure, BMI, cardiovascular fitness, atopy, asthma, infections and lifestyle traits. Blood samples for DNA, lipids, glucose, insulin and hormones were also taken. Polygenic risk scores were calculated for each of the 5404 individuals using their genome-wide data and 48 publicly available GWAS (see **S1 Table**). These were tested for their association with 143 traits measured in the NFBC1966 (see **S2 Table**). Subject overlap between the GWAS summary statistics and the NFBC1966 would cause inflation of the results and so the contribution of the NFBC1966 to the discovery GWAS was either: (1) removed directly by reanalyzing the discovery GWAS with the NFBC1966 data excluded, or if this was not an option then, (2) by recalculating what the meta-analysis GWAS results would have been had the NFBC1966 data been removed based on the effect size estimate and corresponding standard error for each SNP based on GWAS (conducted in the same way as the meta-analysis GWAS) performed on the NFBC1966 only. The latter was enabled via the inverse-variance meta-analyses formulae (this recalculation is approximate if the less commonly used sample-size weighted meta-analysis method was used in the discovery GWAS) as follows:

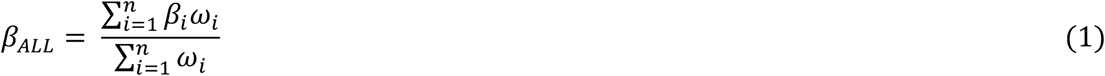

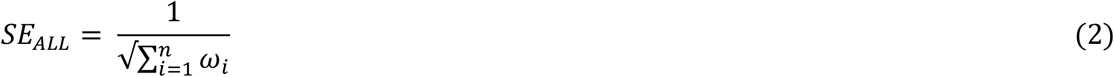

where *β*_*ALL*_ is the effect size estimate meta-analysed across *n* cohorts, where the *β*_*i*_ for each cohort *i* is weighted by the inverse variance of its standard error such that its weight, *ω*_*i*_, equals 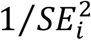.

So, to find the effect size estimate, *β*_*ADJ*_, and *SE*_*ADJ*_,, with cohort *k* removed, by definition:

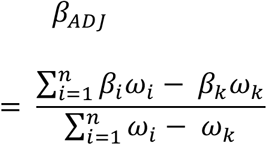

From (1) and (2):

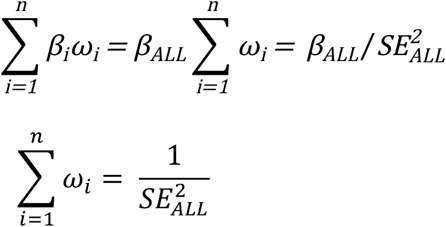

Thus:

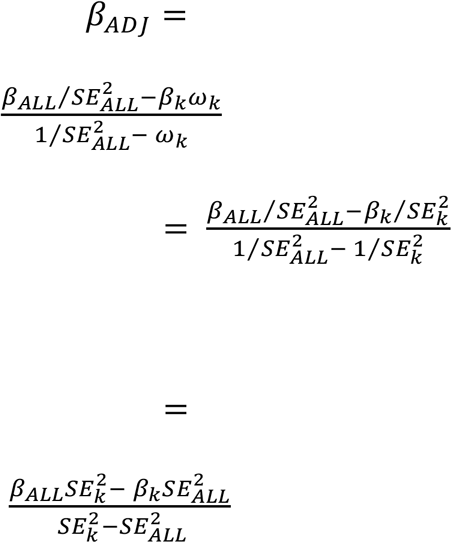

and by definition and (2):

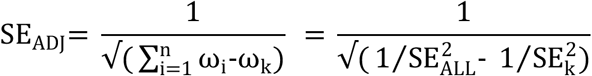

The polygenic risk score software PRSice [13] was used for the data analysis, which involved performing linear (continuous traits) or logistic (binary traits) regression of NFBC1966 target phenotypes on PRS, with PRS computed in relation to each of the 48 GWAS summary statistic results, to test for their association. The SNPs in the base (discovery GWAS) had ambiguous SNP genotype calls removed and strands flipped where necessary. SNPs in linkage disequilibrium (LD) were “clumped” using a threshold of *r*^*2*^ < 0.1 across 250kb windows to ensure that those analysed are largely independent [13]. Ancestry informative covariates were generated from the target genotype data using Principal Component Analysis (PCA), and the first 10 PCs were included in the regressions to control for population stratification. Further analyses also controlled for sex, socio-economic status and BMI to potentially increase power or expose mediation effects (e.g. see **S3 Fig**, **S4 Fig**, **S5 Fig**, **S6 Fig** and **S7 Fig**). High-resolution scoring was performed in PRSice to identify the most predictive PRS for each trait from the large number of PRS that can be formed by inclusion of groups of SNPs with different GWAS association *P*-values thresholds [13]. While this makes the *P*-value for association between the PRS and target phenotypes over-fit, we apply a significance threshold of *P* = 0.004 for each association test based on a permutation study for high-resolution scoring by Euesden et al. [13]. Bonferroni correction for the large number of genetic-phenotype tests performed (6864) produces a conservative significance threshold of *P* < 5×10^−7^ (0.004/6864), with thresholds of *P* < 0.001 and *P* < 0.01 used to indicate potential associations in the data.

Traits pertaining to socio-economic status and exercise measures were originally coded with the highest number on the questionnaire pertaining to the lowest measure of each trait (e.g. “5” for the lowest ability for “running 5km”; see **S3 Table**), so these were recoded in the opposite direction to provide greater clarity in the results.

PRS-sex interactions were also tested for the 162 genetic-phenotype associations that exceeded the significance threshold (P < 5×10^−7^) in linear models with the corresponding outcome trait regressed on PRS, sex, 10 PCs and PRS*sex interaction term. Bonferroni correction for the number of interactions tested produces a conservative significance threshold of *P* < 3×10^−4^ (0.05/162).

## Results

PRS were calculated for 5404 participants with genotype data available in NFBC1966 using publicly available GWAS summary statistics (see **S1 Table**). The PRS across the sample were tested for associations with phenotypes from data collected both in early life and at 31 years in the NFBC1966 participants. Data included anthropometric measurements, blood measurements (e.g. cardiometabolic risk factors), hormone levels, and questionnaire data at baseline and 31 years, socio-economic factors, medical history and health related behaviours. Details of all phenotypes are provided in **S3 Table**. Altogether, PRS were computed using 48 GWAS meta-analysis summary statistics and were tested for their association with 143 NFBC1966 phenotypes, corresponding to 6864 tests in total.

We grouped the results into 6 categories of related target phenotypes: medical conditions (Fig 1), metabolic traits (Fig 2), lifestyle and social factors (Fig 3), health (**S1 Fig**), and anthropometrics (**S2 Fig**). We present the results corresponding to each category by heat maps depicting the associations between each PRS and each target phenotype, with significant associations highlighted by asterisks.

**Fig 1.**
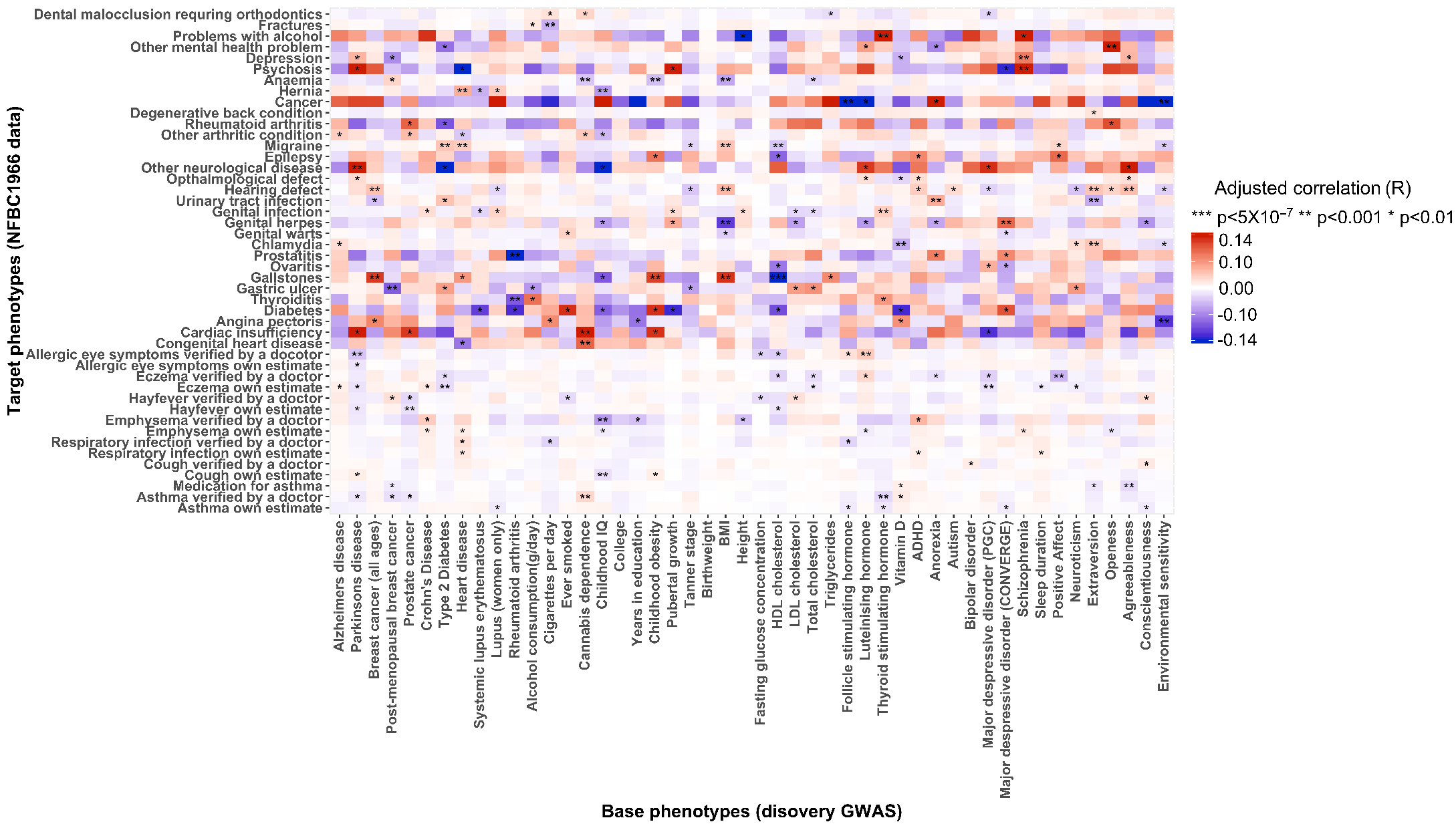
Heat map showing associations between polygenic risk scores from GWAS traits (X-axis) and NFBC1966 traits (Y-axis) for self-reported diseases, medical and psychiatric conditions, verified or treated by a doctor. Asterisks denote different levels of evidence according to *P*-value: *** = *P* < 5×10^−7^, ** = *P* < 0.001, * = *P* < 0.01. Red *r*values indicate positive correlations, while blue *r* values indicate negative correlations.

**Fig 2.**
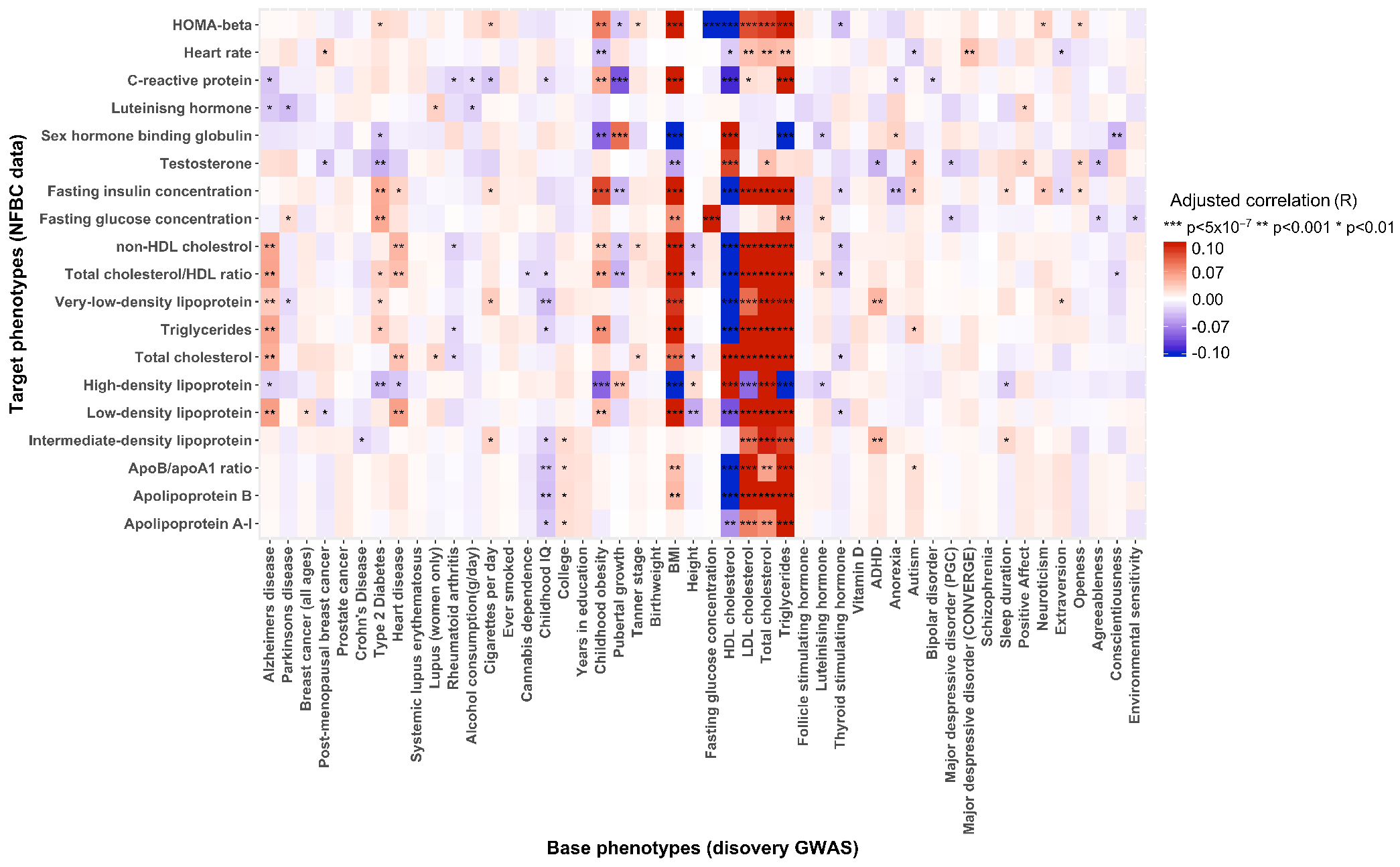
Heat map showing genetic associations between polygenic risk scores of GWAS traits (X-axis) and NFBC1966 traits (Y-axis) for cardiometabolic measurements from blood samples. Asterisks denote different levels of evidence according to *P*-value: *** = *P* < 5×10^−7^, ** = *P* < 0.001, * = *P* < 0.01. Red *r* values indicate positive correlations, while blue *r* values indicate negative correlations.

Given the unusually large number of results that an analysis of this kind produces, and its relative novelty, we recommend considering the following points while inspecting the results: (i) the statistical power is a function of the sample sizes of both the discovery GWAS and target data, and while we have removed those that are highly underpowered there remains high variation in power among the results, (ii) the observation of genetic associations limits the opportunity for confounding to produce spurious associations but the results here do not distinguish between two main plausible explanations for associations: horizontal pleiotropy (genetic effect is on the two traits directly) and vertical pleiotropy (genetic effect is on one trait, which has a downstream effect on the second trait) [2] and the order of any causation is not necessarily from discovery to target trait (see **Discussion**), (iii) the same basic adjustment for covariates (see **Methods**) is performed across all tests, so results may change qualitatively with adjustment of risk factors particularly relevant to the target trait under study, (iv) given the lack of mechanistic insight and replication in these results, they should be viewed more as hypothesis-generating than confirmatory. Our hope is that particular results will motivate and guide follow-up investigations by researchers with expertise in the corresponding phenotypes. The results that we highlight and summarise below reflect only those that we consider some of the more interesting results and are necessarily only a subset of the potentially important results.

Fig 1 shows associations between the 48 PRS and 46 NFBC1966 medical conditions that were either self-estimated by the participant or self-reported as being verified by a physician; these include metabolic disorders, psychiatric disorders, infectious diseases and allergies. None of the associations in this category were significant after applying the stringent Bonferroni correction for multiple testing, which may reflect the small number of cases of many of the conditions in the sample. However, there were a number results with suggestive evidence (P < 0.001) that confirm expectations or that may of interest for follow-up studies. For example, the associations between schizophrenia PRS and depression, schizophrenia and psychosis are as expected, as is that between Parkinson’s disease PRS and self-reported neurological disease [14–16]. The positive association between heart disease PRS and migraine is supported by epidemiological research [17,18], but is in contrast with previous genetic studies suggesting that the genetic component for migraine with aura is protective for heart disease [19]. Moreover, the PRS for HDL cholesterol, a proposed protective factor for cardiovascular diseases [20], was negatively associated with gallstones [21], follicle stimulating hormone with self-reported cancer [22], and thyroid stimulating hormone with asthma [23]. The Cannabis smoking PRS was positively associated with asthma, inborn heart disease and cardiac insufficiency [24,25]. There are also several suggestive associations relating to psychological trait PRS. Positive affect PRS is positively associated with reduced eczema [26], extraversion PRS is positively associated with chlamydia, supporting literature linking extraversion to high risk sexual behaviour [27], the openness PRS is positively associated with mental health problems [28–30], while the PRS on environmental sensitivity is negatively correlated with angina and cancer [31,32].

Fig 2 depicts the associations between the 48 PRS and 19 NFBC1966 cardiometabolic traits. There are a large number of significant results here, including many that would be expected based on the epidemiological literature. Some associations have been observed in the genetic literature previously, such as between BMI PRS and the major lipids (LDL, HDL, cholesterol) and C-reactive protein [33], while others are novel, such as between the lipids and testosterone, sex hormone binding globulin and insulin [34]. A genetic overlap between Alzheimer’s disease and plasma lipids HDL, LDL and triglycerides was replicated [35], but there were additional associations with total cholesterol and VLDL here. VLDL also shows suggestive evidence of associations with the PRS of childhood IQ, ADHD and cigarette smoking, indicating a potentially greater role for this lipoprotein than previously thought [38,39]. The associations between cardiovascular disease and diabetes PRS and the lipids are as expected from the epidemiological literature [36,37], while the suggestive evidence for associations between anorexia and neuroticism PRS and insulin support the proposed role for genetics in the shared aetiology between insulin and cognitive function [40].

Fig 3 shows the associations between the 48 PRS and 38 NFBC1966 lifestyle and social factors, mostly comprising occupation, smoking and alcohol consumption measures. The results pertaining to education highlight the potential for mediation by lifestyle to produce genetic pleiotropy; for example, the college and ‘years in education’ PRS are negatively associated with beer/cider and wine amount but positively associated with wine frequency, which may reflect the adoption of different social lifestyles according to attendance at university and adulthood socio-economic position. This is supported by the associations between the education PRS and socio-economic status measures here, and also by twin studies linking education and health behaviours [41, 42]. The PRS for HDL is positively correlated with most of the alcohol consumption measures and negatively correlated with the smoking measures, while the opposite pattern is observed for the Triglycerides PRS. However, both the HDL and Triglycerides PRS have strong positive associations with the Oral Contraceptive Pill (OCP), which most likely reflects the impact of increased lipid levels among individuals on OCP in the lipids GWAS samples; that is, genetic factors affecting uptake of OCP may have been captured by the lipids GWAS due to the lipid-altering effect of OCP. Birth weight PRS was positively associated with both mother’s age and father’s age. This reflects findings in the literature linking lower maternal age with increased odds of low birth weights, although the association is U-shaped [43]. It has been suggested that social disadvantage underlies the low maternal age-low birth weight link [44]; nevertheless, our data suggest that whatever the underlying causal factor is, it is under a degree of genetic control.

**Fig 3.**
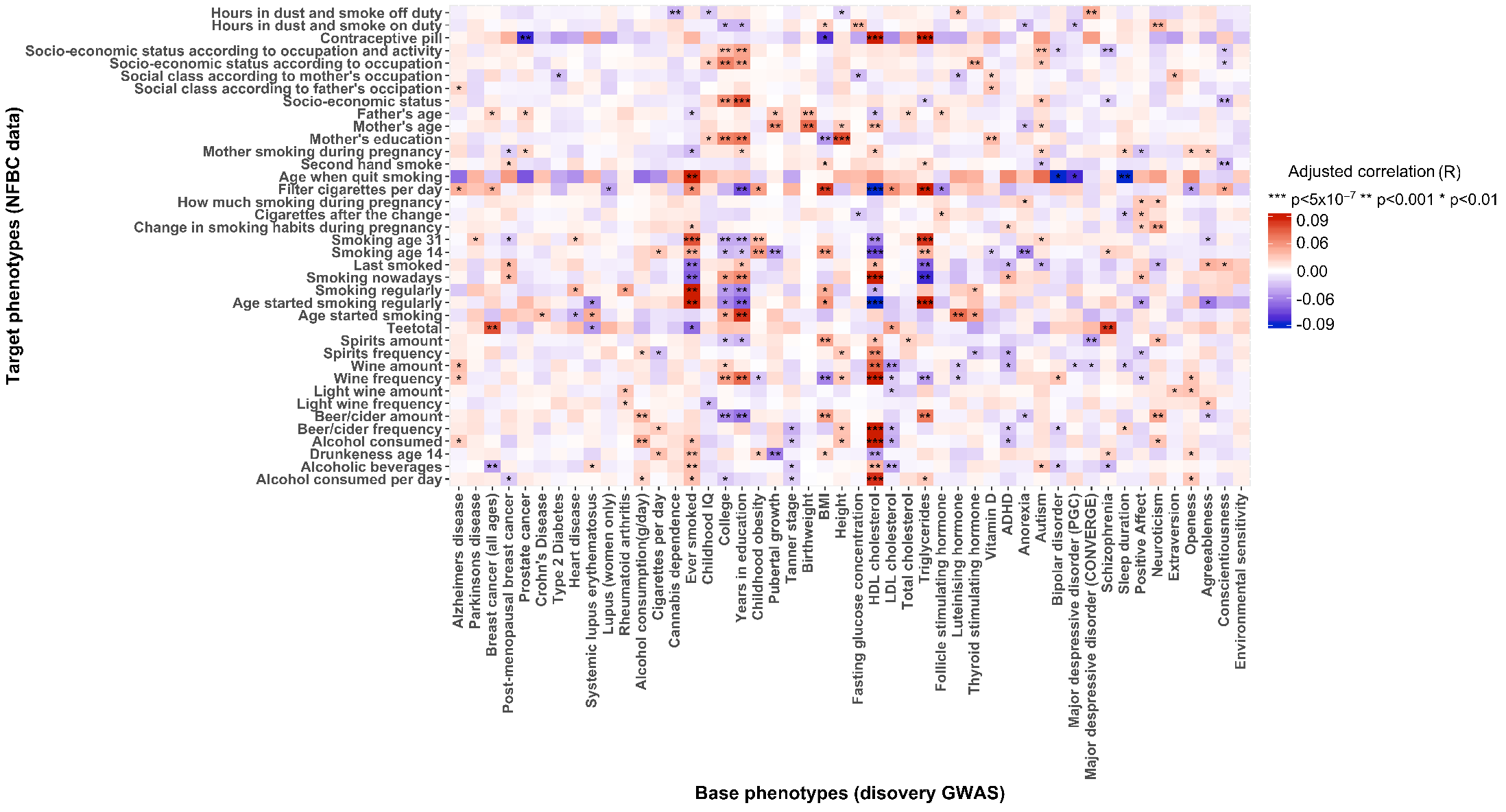
Heat map showing genetic associations between polygenic risk scores of GWAS traits (X-axis) and NFBC1966 traits (Y-axis) from questionnaires on lifestyle and social factors. Asterisks denote different levels of evidence according to *P*-value: *** = *P* < 5×10^−7^, ** = *P* < 0.001, * = *P* < 0.01. Red *r* values indicate positive correlations, while blue *r* values indicate negative correlations.

**S1 Fig** and **S2 Fig** display the associations between the 48 PRS and anthropometrics and health traits. The associations relating to traits such as height, weight, blood pressure, physical activity and diabetes are as expected based on the literature [45–47], but the heat maps also reveal potentially novel insights. For example, exercise, especially running, is positively correlated with the PRS for childhood IQ, years in education, bipolar disorder and positive affect, but negatively correlated with the ADHD PRS, which may highlight a potentially important, but mixed, role of physical activity in psychiatric disorders. Exercise has been shown to alleviate psychiatric symptoms [48,49] but since there are relatively few individuals in our data with psychiatric disorders then these associations may be more likely due to mediation by exercise rather than its therapeutic effects. Likewise, the PRS for breast cancer, Crohn’s disease and diabetes are associated with several physical activity measures as expected from epidemiological findings [50,51].

## PRS*Sex interactions

Interactions between PRS and sex were also investigated (see **Methods**). The top results from the interaction analyses are presented in Fig 4. A significant effect modification by sex of the association between HDL PRS and sex hormone binding globulin levels (*P* = 8.13×10^−6^) was observed, while several other interactions were only marginally significant (*P* < 0.05). These results reflect the general finding in the literature that the autosomal genetic influence of complex traits is largely similar between males and females, with genotype by sex interactions having very small effect sizes compared to the main effects [52].

**Fig 4.**
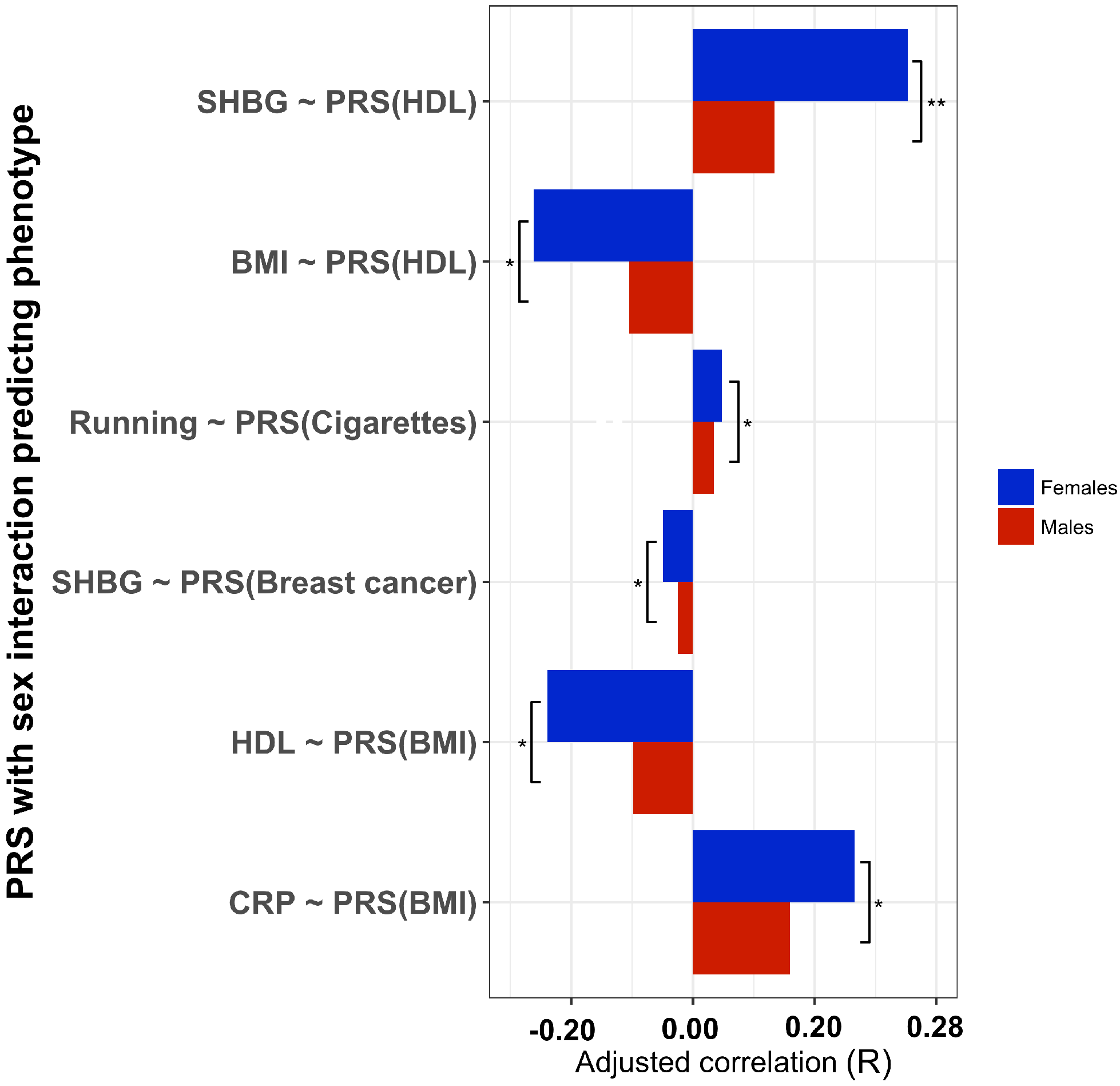
Bar plot showing top-ranking sex interactions with *r* values for males and females. Y-axis shows phenotype predicted in regression with predicting PRS in parenthesis. Asterisks denote evidence for association in terms of *P*-value: ** = *P* < 3×10^−4^, * = *P* < 0.05. Full results in **S2 table**.

## Discussion

Here we performed a large-scale systematic survey of genetic-phenotype associations using a set of 48 GWAS summary statistics and 143 phenotypes measured in the Northern Finland Birth Cohort 1966 (NFBC1966). Novel associations and replications were identified across a broad array of clinical, cardiometabolic, anthropometric, infectious disease, psychiatric and lifestyle traits. While this study is among a growing number of large cross-trait studies investigating shared genetic aetiology among human phenotypes, it represents the largest medically focused such study using the polygenic risk score (PRS) approach to date [6,7]. The use of the PRS method here highlights the potential for exploiting a single epidemiological study to gain insights into the underlying aetiology of a huge number of phenotypes, given the rich phenotyping typical of such studies. This is in contrast to the popular LD Score regression method [4], which requires large GWAS to have been performed on all traits under study and does not allow control for covariates.

The large number of GWAS exploited here and depth of phenotyping of the NFBC1966 meant that patterns of genetic-phenotype associations corresponding to related traits emerged, offering both internal support for associations as well as highlighting apparently conflicting results that deserve specific follow-up. For instance, associations between education PRS and a range of alcohol and smoking measures that potentially indicate mediation via socio-economic position, are supported by associations between the education PRS and socio-economic status variables. However, while the PRS for HDL and Triglycerides had associations in the opposite direction across almost all target traits, they had the same strong positive correlation with the oral contraceptive pill. Therefore, a profile of associations is observed among related traits, particularly useful for highlighting potential causal pathways and guiding follow-up investigations.

The central limitation of such a large-scale systematic study is that the testing performed on any specific phenotype is inevitably superficial in nature. While some of the genetic-phenotype associations observed suggest particular aetiological explanations, especially when considering groups of related associations, other than sex-interaction analyses we performed no further statistical testing to gain additional insights. However, we believe that this large-scale hypothesis-free approach to investigating shared genetic aetiology among human phenotypes has much value: within a single consistent analysis we have revealed evidence for shared genetic aetiology among hundreds of traits and provided what may be considered a ‘treasure trove’ of avenues for follow-up investigations. As similar studies are performed on different data sets and populations, patterns of replicating associations will emerge. The possibility of ‘collider bias’ [53] should be considered in relation to any of the observed associations but should be minimized by the use of a birth cohort, in which the vast majority of births in Northern Finland in 1966 were included and a high proportion of these genotyped. While spurious pleiotropy is also possible [2], the fact that these are genetic associations should be a greater indication of genuine causation than classical epidemiological associations. For instance, the associations observed here between alcohol consumption, smoking, education and socio-economic status suggest causal links between these factors, which has implications for epidemiology, in which these measures are often considered as only confounded by each other. However, such inference relating to these associations is only speculative until rigorous follow-up investigations are performed to uncover causal mechanisms.

This study has demonstrated that taking a hypothesis-free polygenic risk score approach to the investigation of shared genetic aetiology among phenotypes is an effective way of replicating previous, and uncovering novel, genetic-phenotype associations. The key advantage of requiring a relatively small target sample is the opportunity to exploit a greater depth of phenotyping, revealing a higher resolution profile of genetic overlap than possible otherwise. Our hope is that these genetic-phenotype associations provide a foundation and guide for investigations to reveal the pathways that lead to disease, both internally within the body, and externally through mediation via behavior and lifestyle.

## Acknowledgements

We thank the late professor Paula Rantakallio (launch of NFBC1966), the participants in the 31yrs study and the NFBC project centre.

We thank the International Genomics of Alzheimer’s Project (IGAP) for providing summary results data for these analyses. The investigators within IGAP contributed to the design and implementation of IGAP and/or provided data but did not participate in analysis or writing of this report. IGAP was made possible by the generous participation of the control subjects, the patients, and their families. The i–Select chips was funded by the French National Foundation on Alzheimer’s disease and related disorders. EADI was supported by the LABEX (laboratory of excellence program investment for the future) DISTALZ grant, Inserm, Institut Pasteur de Lille, Université de Lille 2 and the Lille University Hospital. GERAD was supported by the Medical Research Council (Grant n° 503480), Alzheimer’s Research UK (Grant n° 503176), the Wellcome Trust (Grant n° 082604/2/07/Z) and German Federal Ministry of Education and Research (BMBF): Competence Network Dementia (CND) grant n° 01GI0102, 01GI0711, 01GI0420. CHARGE was partly supported by the NIH/NIA grant R01 AG033193 and the NIA AG081220 and AGES contract N01–AG–12100, the NHLBI grant R01 HL105756, the Icelandic Heart Association, and the Erasmus Medical Center and Erasmus University. ADGC was supported by the NIH/NIA grants: U01 AG032984, U24 AG021886, U01 AG016976, and the Alzheimer’s Association grant ADGC–10–196728. P.F.O receives funding from the UK Medical Research Council (MR/N015746/1) and the Wellcome Trust (109863/Z/15/Z). This report represents independent research (part)-funded by the National Institute for Health Research (NIHR) Biomedical Research Centre at South London and Maudsley NHS Foundation Trust and King’s College London. The views expressed are those of the authors and not necessarily those of the NHS, the NIHR, or the Department of Health. Data on coronary artery disease / myocardial infarction have been contributed by CARDIoGRAMplusC4D investigators and have been downloaded from http://www.CARDIOGRAMPLUSC4D.ORG.

Data on head circumference, childhood obesity, pubertal growth, tanner stage and birth weight traits has been contributed by EGG Consortium and has been downloaded from http://www.egg-consortium.org.

The Educational Attainment GWAS results were accessed under the Data Sharing Agreement of the Social Science Genetic Association Consortium. We thank the SSGAC for facilitating this research. C.A.R. acknowledges funding from the Netherlands Organisation for Scientific Research (NWO Veni grant 016.165.004)

## References

1. Sivakumaran S, Agakov F, Theodoratou E, Prendergast JG, Zgaga L, Manolio T, Rudan I, McKeigue P, Wilson JF, Campbell H. Abundant pleiotropy in human complex diseases and traits. American Journal of Human Genetics. 2011 Nov 11;89(5):607–18.

2. Solovieff N, Cotsapas C, Lee PH, Purcell SM, Smoller JW. Pleiotropy in complex traits: challenges and strategies. Nature Reviews Genetics. 2013 Jul 1;14(7):483–95.

3. Trzaskowski M, Yang J, Visscher PM, Plomin R. DNA evidence for strong genetic stability and increasing heritability of intelligence from age 7 to 12. Molecular Psychiatry. 2014 Mar 1;19(3):380–4.

4. Bulik-Sullivan BK, Loh PR, Finucane HK, Ripke S, Yang J, Patterson N, Daly MJ, Price AL, Neale BM, Schizophrenia Working Group of the Psychiatric Genomics Consortium. LD Score regression distinguishes confounding from polygenicity in genome-wide association studies. Nature Genetics. 2015 Mar 1;47(3):291–5.

5. Smoller JW, Kendler K, Craddock NJ, Lee PH, Neale BM, Nurnberger JI, Ripke S, Santangelo S, Sullivan PF, Purcell S, Anney R. Identification of risk loci with shared effects on five major psychiatric disorders: a genome-wide analysis. Lancet. 2013 Apr 20;381(9875):1371–9.

6. Bulik-Sullivan B, Finucane HK, Anttila V, Gusev A, Day FR, Loh PR, Duncan L, Perry JR, Patterson N, Robinson EB, Daly MJ. An atlas of genetic correlations across human diseases and traits. Nature Genetics. 2015 Sep 28.

7. Krapohl E, Euesden J, Zabaneh D, Pingault JB, Rimfeld K, Von Stumm S, Dale PS, Breen G, O’reilly PF, Plomin R. Phenome-wide analysis of genome-wide polygenic scores. Molecular Psychiatry. 2016 Sep 1;21(9):1188–93.

8. Hagenaars SP, Harris SE, Davies G, Hill WD, Liewald DC, Ritchie SJ, Marioni RE, Fawns-Ritchie C, Cullen B, Malik R, Worrall BB. Shared genetic aetiology between cognitive functions and physical and mental health in UK Biobank (N= 112 151) and 24 GWAS consortia. Molecular Psychiatry. 2016 Nov 1;21(11):1624–32.

9. Chatterjee N, Shi J, Garda-Closas M. Developing and evaluating polygenic risk prediction models for stratified disease prevention. Nature Reviews Genetics. 2016 Jul 1;17(7):392–406.

10. Breen G, Li Q, Roth BL, O’Donnell P, Didriksen M, Dolmetsch R, O’Reilly PF, Gaspar HA, Manji H, Huebel C, Kelsoe JR. Translating genome-wide association findings into new therapeutics for psychiatry. Nature Neuroscience. 2016 Nov 1;19(11):1392–6.

11. Dudbridge F. Polygenic epidemiology. Genetic epidemiology. 2016 May 1;40(4):268–72.

12. Sorri M, Järvelin MR. Well-being and health. Background to the northern Finland 1966 birth cohort research. International Journal of Circumpolar Health. 1998 Jul 1;57(2-3):82.

13. Euesden J, Lewis CM, O’Reilly PF. PRSice: polygenic risk score software. Bioinformatics. 2015 May 1;31(9):1466–8.

14. Schulze TG, Akula N, Breuer R, Steele J, Nalls MA, Singleton AB, Degenhardt FA, Nöthen MM, Cichon S, Rietschel M, Mcmahon FJ. Molecular genetic overlap in bipolar disorder, schizophrenia, and major depressive disorder. The World Journal of Biological Psychiatry. 2014 Apr 1;15(3):200–8.

15. Ivleva E, Thaker G, Tamminga CA. Comparing genes and phenomenology in the major psychoses: schizophrenia and bipolar 1 disorder. Schizophrenia Bulletin. 2008 Jul 1;34(4):734–42.

16. Lewis PA, Cookson MR. Gene expression in the Parkinson’s disease brain. Brain Research Bulletin. 2012 Jul 1;88(4):302–12.

17. Sacco S, Ornello R, Ripa P, Tiseo C, Degan D, Pistoia F, Carolei A. Migraine and risk of ischaemic heart disease: a systematic review and meta-analysis of observational studies. European Journal of Neurology. 2015 Jun 1; 22(6):1001–11

18. Bigal ME, Kurth T, Hu H, Santanello N, Lipton RB. Migraine and cardiovascular disease Possible mechanisms of interaction. Neurology. 2009 May 26;72(21):1864–71.

19. Winsvold BS, Nelson CP, Malik R, Gormley P, Anttila V, Vander Heiden J, Elliott KS, Jacobsen LM, Palta P, Amin N, de Vries B. Genetic analysis for a shared biological basis between migraine and coronary artery disease. Neurology Genetics. 2015 Jun 1;1(1):e10.

20. Ali KM, Wonnerth A, Huber K, Wojta J. Cardiovascular disease risk reduction by raising HDL cholesterol-current therapies and future opportunities. British journal of pharmacology. 2012 Nov 1;167(6):1177–94.

21. Tîrziu S, Bel S, Bondor CI, Acalovschi M. Risk factors for gallstone disease in patients with gallstones having gallstone heredity. A case-control study. Romanian Journal of Internal Medicine= Revue Roumaine de Médecine Interne. 2008;46(3):223.

22. McSorley MA, Alberg AJ, Allen DS, Allen NE, Brinton LA, Dorgan JF, Kaaks R, Rinaldi S, Helzlsouer KJ. Prediagnostic circulating follicle stimulating hormone concentrations and ovarian cancer risk. International Journal of Cancer. 2009 Aug 1;125(3):674–9.

23. Oppedal RJ, Khan DA, Brown ES. Hypothyroidism in patients with asthma and major depressive disorder. Primary care companion to the Journal of clinical psychiatry. 2007;9(6):467.

24. Tetrault JM, Crothers K, Moore BA, Mehra R, Concato J, Fiellin DA. Effects of marijuana smoking on pulmonary function and respiratory complications: a systematic review. Archives of internal medicine. 2007 Feb 12;167(3):221–8.

25. Mukamal KJ, Maclure M, Muller JE, Mittleman MA. An exploratory prospective study of marijuana use and mortality following acute myocardial infarction. American Heart Journal. 2008 Mar 31;155(3):465–70.

26. Schmitt J, Romanos M, Pfennig A, Leopold K, Meurer M. Psychiatric comorbidity in adult eczema. British Journal of Dermatology. 2009 Oct 1;161(4):878–83.

27. Miller JD, Lynam D, Zimmerman RS, Logan TK, Leukefeld C, Clayton R. The utility of the Five Factor Model in understanding risky sexual behavior. Personality and Individual Differences. 2004 May 31;36(7):1611–26.

28. Kotov R, Gamez W, Schmidt F, Watson D. Linking “big” personality traits to anxiety, depressive, and substance use disorders: a meta-analysis. Psychological Bulletin. 2010 Sep;136(5):768.

29. Lamers SM, Westerhof GJ, Kovács V, Bohlmeijer ET. Differential relationships in the association of the Big Five personality traits with positive mental health and psychopathology. Journal of Research in Personality. 2012 Oct 31;46(5):517–24.

30. Widiger TA, Trull TJ. Personality and psychopathology: an application of the Five-Factor Model. Journal of Personality. 1992 Jun 1;60(2):363–93.

31. Friedman HS, Booth-Kewley S. The “disease-prone personality”: A meta-analytic view of the construct. American Psychologist. 1987 Jun;42(6):539.

32. Chida Y, Hamer M, Wardle J, Steptoe A. Do stress-related psychosocial factors contribute to cancer incidence and survival? Nature Clinical Practice Oncology. 2008 Aug 1;5(8):466–75.

33. Andreassen OA, Desikan RS, Wang Y, Thompson WK, Schork AJ, Zuber V, Doncheva NT, Ellinghaus E, Albrecht M, Mattingsdal M, Franke A. Abundant genetic overlap between blood lipids and immune-mediated diseases indicates shared molecular genetic mechanisms. PLoS One. 2015 Apr 8;10(4):e0123057.

34. Zhang N, Zhang H, Zhang X, Zhang B, Wang F, Wang C, Zhao M, Yu C, Gao L, Zhao J, Guan Q. The relationship between endogenous testosterone and lipid profile in middle-aged and elderly Chinese men. European Journal of Endocrinology. 2014 Apr 1;170(4):487–94.

35. Desikan RS, Schork AJ, Wang Y, Thompson WK, Dehghan A, Ridker PM, Chasman DI, McEvoy LK, Holland D, Chen CH, Karow DS. Polygenic Overlap Between C-Reactive Protein, Plasma Lipids, and Alzheimer DiseaseCLINICAL PERSPECTIVE. Circulation. 2015 Jun 9;131(23):2061–9.

36. Saxena R, Voight BF, Lyssenko V, Burtt NP, de Bakker PI, Chen H, Roix JJ, Kathiresan S, Hirschhorn JN, Daly MJ, Hughes TE. Genome-wide association analysis identifies loci for type 2 diabetes and triglyceride levels. Science. 2007 Jun 1;316(5829):1331–6.

37. Aulchenko YS, Ripatti S, Lindqvist I, Boomsma D, Heid IM, Pramstaller PP, Penninx BW, Janssens AC, Wilson JF, Spector T, Martin NG. Loci influencing lipid levels and coronary heart disease risk in 16 European population cohorts. Nature Genetics. 2009 Jan 1;41(1):47–55.

38. Taylor AE, Guthrie PA, Smith GD, Golding J, Sattar N, Hingorani AD, Deanfield JE, Day IN. IQ, educational attainment, memory and plasma lipids: associations with apolipoprotein E genotype in 5995 children. Biological Psychiatry. 2011 Jul 15;70(2):152–8.

39. Greenfield JR, Samaras K, Jenkins AB, Kelly PJ, Spector TD, Campbell LV. Do gene–environment interactions influence fasting plasma lipids? A study of twins. European Journal of Clinical Investigation. 2004 Sep 1;34(9):590–8.

40. Strachan MW. Insulin and cognitive function in humans: experimental data and therapeutic considerations. Biochemical Society Transactions. 2005 Nov 1;33(Pt 5):1037.

41. Johnson W, Kyvik KO, Mortensen EL, Skytthe A, Batty GD, Deary IJ. Does education confer a culture of healthy behavior? Smoking and drinking patterns in Danish twins. American Journal of Epidemiology. 2010 Nov 4:kwq333.

42. Benner AD, Kretsch N, Harden KP, Crosnoe R. Academic achievement as a moderator of genetic influences on alcohol use in adolescence. Developmental Psychology. 2014 Apr;50(4):1170.

43. Restrepo-Méndez MC, Lawlor DA, Horta BL, Matijasevich A, Santos IS, Menezes A, Barros FC, Victora CG. The association of maternal age with birth weight and gestational age: a cross-cohort comparison. Paediatric and Perinatal Epidemiology. 2015 Jan 1;29(1):31–40.

44. Dennis JA, Mollborn S. Young maternal age and low birth weight risk: an exploration of racial/ethnic disparities in the birth outcomes of mothers in the United States. The Social Science Journal. 2013 Dec 31;50(4):625–34.

45. Hemmingsson E, Ekelund U. Is the association between physical activity and body mass index obesity dependent? International Journal of Obesity. 2007 Apr 1;31(4):663–8.

46. Carlsson S, Ahlbom A, Lichtenstein P, Andersson T. Shared genetic influence of BMI, physical activity and type 2 diabetes: a twin study. Diabetologia. 2013 May 1;56(5):1031–5.

47. Dubois L, Kyvik KO, Girard M, Tatone-Tokuda F, Pérusse D, Hjelmborg J, Skytthe A, Rasmussen F, Wright MJ, Lichtenstein P, Martin NG. Genetic and environmental contributions to weight, height, and BMI from birth to 19 years of age: an international study of over 12,000 twin pairs. PLOS One. 2012 Feb 8;7(2):e30153.

48. Archer T, Kostrzewa RM. Physical exercise alleviates ADHD symptoms: regional deficits and development trajectory. Neurotoxicity Research. 2012 Feb 1;21(2):195–209.

49. Thomson D, Turner A, Lauder S, Gigler ME, Berk L, Singh AB, Pasco JA, Berk M, Sylvia L. A brief review of exercise, bipolar disorder, and mechanistic pathways. Frontiers in Psychology. 2015;6.

50. Warburton DE, Nicol CW, Bredin SS. Health benefits of physical activity: the evidence. Canadian Medical Association Journal. 2006 Mar 14;174(6):801–9.

51. Ng V, Millard W, Lebrun C, Howard J. Exercise and Crohn’s disease: speculations on potential benefits. Canadian Journal of Gastroenterology and Hepatology. 2006;20(10):657–60.

52. Kassam I, McRae AF. The autosomal genetic control of sexually dimorphic traits in humans is largely the same across the sexes. Genome Biology. 2016 Aug 5;17(1):169.

53. Munafò MR, Tilling K, Taylor AE, Evans DM, Davey Smith G. Collider scope: when selection bias can substantially influence observed associations. International Journal of Epidemiology. 2017 Sep 27: dyx206

54. Lambert JC, Ibrahim-Verbaas CA, Harold D, Naj AC, Sims R, Bellenguez C, Jun G, DeStefano AL, Bis JC, Beecham GW, Grenier-Boley B. Meta-analysis of 74,046 individuals identifies 11 new susceptibility loci for Alzheimer’s disease. Nature Genetics. 2013 Dec 1;45(12):1452–8.

55. Plagnol V, Nalls MA, Bras JM, Hernandez DG, Sharma M, Sheerin UM, Saad M, Simon-Sanchez J, Schulte C, Lesage S, Sveinbjornsdottir S. A Two-Stage Meta-Analysis Identifies Several New Loci for Parkinson’s Disease. PLoS Genetics. 2011;7(6).

56. Murabito JM, Rosenberg CL, Finger D, Kreger BE, Levy D, Splansky GL, Antman K, Hwang SJ. A genome-wide association study of breast and prostate cancer in the NHLBI’s Framingham Heart Study. BMC Medical Genetics. 2007 Sep 19;8(1):S6.

57. Hunter DJ, Kraft P, Jacobs KB, Cox DG, Yeager M, Hankinson SE, Wacholder S, Wang Z, Welch R, Hutchinson A, Wang J. A genome-wide association study identifies alleles in FGFR2 associated with risk of sporadic postmenopausal breast cancer. Nature Genetics. 2007 Jul 1;39(7):870–4.

58. Jostins L, Ripke S, Weersma RK, Duerr RH, McGovern DP, Hui KY, Lee JC, Schumm LP, Sharma Y, Anderson CA, Essers J. Host-microbe interactions have shaped the genetic architecture of inflammatory bowel disease. Nature. 2012 Nov 1;491(7422):119–24.

59. Morris AP, Voight BF, Teslovich TM, Ferreira T, Segre AV, Steinthorsdottir V, Strawbridge RJ, Khan H, Grallert H, Mahajan A, Prokopenko I. Large-scale association analysis provides insights into the genetic architecture and pathophysiology of type 2 diabetes. Nature Genetics. 2012 Sep;44(9):981.

60. Schunkert H, König IR, Kathiresan S, Reilly MP, Assimes TL, Holm H, Preuss M, Stewart AF, Barbalic M, Gieger C, Absher D. Large-scale association analysis identifies 13 new susceptibility loci for coronary artery disease. Nature Genetics. 2011 Apr 1;43(4):333–8.

61. Hom G, Graham RR, Modrek B, Taylor KE, Ortmann W, Garnier S, Lee AT, Chung SA, Ferreira RC, Pant PK, Ballinger DG. Association of systemic lupus erythematosus with C8orf13–BLK and ITGAM–ITGAX. New England Journal of Medicine. 2008 Feb 28;358(9):900–9.

62. Ramos PS, Criswell LA, Moser KL, Comeau ME, Williams AH, Pajewski NM, Chung SA, Graham RR, Zidovetzki R, Kelly JA, Kaufman KM. A comprehensive analysis of shared loci between systemic lupus erythematosus (SLE) and sixteen autoimmune diseases reveals limited genetic overlap. PLoS Genetics. 2011 Dec 8;7(12):e1002406.

63. Chen R, Stahl EA, Kurreeman FA, Gregersen PK, Siminovitch KA, Worthington J, Padyukov L, Raychaudhuri S, Plenge RM. Fine mapping the TAGAP risk locus in rheumatoid arthritis. Genes and Immunity. 2011 Jun 1;12(4):314–8.

64. Schumann G, Liu C, O’Reilly P, Gao H, Song P, Xu B, Ruggeri B, Amin N, Jia T, Preis S, Lepe MS. KLB is associated with alcohol drinking, and its gene product β-Klotho is necessary for FGF21 regulation of alcohol preference. Proceedings of the National Academy of Sciences. 2016 Nov 28:201611243.

65. Tobacco and Genetics Consortium. Genome-wide meta-analyses identify multiple loci associated with smoking behavior. Nature Genetics. 2010 May 1;42(5):441–7.

66. Verweij KJ, Vinkhuyzen AA, Benyamin B, Lynskey MT, Quaye L, Agrawal A, Gordon SD, Montgomery GW, Madden P, Heath AC, Spector TD. The genetic aetiology of cannabis use initiation: a meta-analysis of genome-wide association studies and a SNP-based heritability estimation. Addiction Biology. 2013 Sep 1;18(5):846–50.

67. Rietveld CA, Esko T, Davies G, Pers TH, Turley P, Benyamin B, Chabris CF, Emilsson V, Johnson AD, Lee JJ, De Leeuw C. Common genetic variants associated with cognitive performance identified using the proxy-phenotype method. Proceedings of the National Academy of Sciences. 2014 Sep 23;111(38):13790–4.

68. Rietveld CA, Medland SE, Derringer J, Yang J, Esko T, Martin NW, Westra HJ, Shakhbazov K, Abdellaoui A, Agrawal A, Albrecht E. GWAS of 126,559 individuals identifies genetic variants associated with educational attainment. Science. 2013 Jun 21;340(6139):1467–71.

69. Bradfield JP, Taal HR, Timpson NJ, Scherag A, Lecoeur C, Warrington NM, Hypponen E, Holst C, Valcarcel B, Thiering E, Salem RM. A genome-wide association meta-analysis identifies new childhood obesity loci. Nature Genetics. 2012 May 1;44(5):526–31.

70. Horikoshi M, Yaghootkar H, Mook-Kanamori DO, Sovio U, Taal HR, Hennig BJ, Bradfield JP, St Pourcain B, Evans DM, Charoen P, Kaakinen M. New loci associated with birth weight identify genetic links between intrauterine growth and adult height and metabolism. Nature Genetics. 2013 Jan 1;45(1):76–82.

71. Cousminer DL, Stergiakouli E, Berry DJ, Ang W, Groen-Blokhuis MM, Körner A, Siitonen N, Ntalla I, Marinelli M, Perry JR, Kettunen J. Genome-wide association study of sexual maturation in males and females highlights a role for body mass and menarche loci in male puberty. Human Molecular Genetics. 2014 Apr 25;23(16):4452–64.

72. Cousminer DL, Berry DJ, Timpson NJ, Ang W, Thiering E, Byrne EM, Taal HR, Huikari V, Bradfield JP, Kerkhof M, Groen-Blokhuis MM. Genome-wide association and longitudinal analyses reveal genetic loci linking pubertal height growth, pubertal timing and childhood adiposity. Human Molecular Genetics. 2013 Feb 27;22(13):2735–47.

73. Locke AE, Kahali B, Berndt SI, Justice AE, Pers TH, Day FR, Powell C, Vedantam S, Buchkovich ML, Yang J, Croteau-Chonka DC. Genetic studies of body mass index yield new insights for obesity biology. Nature. 2015 Feb 12;518(7538):197–206.

74. Allen HL, Estrada K, Lettre G, Berndt SI, Weedon MN, Rivadeneira F, Willer CJ, Jackson AU, Vedantam S, Raychaudhuri S, Ferreira T. Hundreds of variants clustered in genomic loci and biological pathways affect human height. Nature. 2010 Oct 14;467(7317):832–8.

75. Manning AK, Hivert MF, Scott RA, Grimsby JL, Bouatia-Naji N, Chen H, Rybin D, Liu CT, Bielak LF, Prokopenko I, Amin N. A genome-wide approach accounting for body mass index identifies genetic variants influencing fasting glycemic traits and insulin resistance. Nature Genetics. 2012 Jun 1;44(6):659–69.

76. Willer CJ, Schmidt EM, Sengupta S, Peloso GM, Gustafsson S, Kanoni S, Ganna A, Chen J, Buchkovich ML, Mora S, Beckmann JS. Discovery and refinement of loci associated with lipid levels. Nature Genetics. 2013 Nov;45(11):1274–85.

77. Hwang SJ, Yang Q, Meigs JB, Pearce EN, Fox CS. A genome-wide association for kidney function and endocrine-related traits in the NHLBI’s Framingham Heart Study. BMC Medical Genetics. 2007 Sep 19;8(1):S10.

78. Wang TJ, Zhang F, Richards JB, Kestenbaum B, Van Meurs JB, Berry D, Kiel DP, Streeten EA, Ohlsson C, Koller DL, Peltonen L. Common genetic determinants of vitamin D insufficiency: a genome-wide association study. Lancet. 2010 Jul 23;376(9736):180–8.

79. Neale BM, Medland SE, Ripke S, Asherson P, Franke B, Lesch KP, Faraone SV, Nguyen TT, Schäfer H, Holmans P, Daly M. Meta-analysis of genome-wide association studies of attention-deficit/hyperactivity disorder. Journal of the American Academy of Child & Adolescent Psychiatry. 2010 Sep 30;49(9):884–97.

80. Boraska V, Franklin CS, Floyd JA, Thornton LM, Huckins LM, Southam L, Rayner NW, Tachmazidou I, Klump KL, Treasure J, Lewis CM. A genome-wide association study of anorexia nervosa. Molecular Psychiatry. 2014 Oct 1;19(10):1085–94.

81. Smoller JW, Kendler K, Craddock NJ, Lee PH, Neale BM, Nurnberger JI, Ripke S, Santangelo S, Sullivan PF, Purcell S, Anney R. Identification of risk loci with shared effects on five major psychiatric disorders: a genome-wide analysis. Lancet. 2013 Apr 20;381(9875):1371–9.

82. Sklar P, Ripke S, Scott LJ, Andreassen OA, Cichon S, Landén M, Craddock N, Edenberg HJ, Nurnberger JI, Rietschel M, Blackwood D. Large-scale genome-wide association analysis of bipolar disorder identifies a new susceptibility locus near ODZ4. Nature Genetics. 2011;43(10):977–U162.

83. Ripke S, Wray NR, Lewis CM, Hamilton SP, Weissman MM, Breen G, Byrne EM, Blackwood DH, Boomsma DI, Cichon S, Heath AC. A mega-analysis of genome-wide association studies for major depressive disorder. Molecular Psychiatry. 2013 Apr 1;18(4):497–511.

84. Cai N, Bigdeli TB, Kretzschmar W, Li Y, Liang J, Song L, Hu J, Li Q, Jin W, Hu Z, Wang G. Sparse whole-genome sequencing identifies two loci for major depressive disorder. Nature. 2015 Jul;523:588–91.

85. Ripke S, O’Dushlaine C, Chambert K, Moran JL, Kähler AK, Akterin S, Bergen SE, Collins AL, Crowley JJ, Fromer M, Kim Y. Genome-wide association analysis identifies 13 new risk loci for schizophrenia. Nature Genetics. 2013 Oct 1;45(10):1150–9.

86. Hart AB, Engelhardt BE, Wardle MC, Sokoloff G, Stephens M, de Wit H, Palmer AA. Genome-wide association study of d-amphetamine response in healthy volunteers identifies putative associations, including cadherin 13 (CDH13). PloS One. 2012 Aug 28;7(8):e42646.

87. Gottlieb DJ, T O’Connor G, Wilk JB. Genome-wide association of sleep and circadian phenotypes. BMC Medical Genetics. 2007 Sep 19;8(1):S9.

88. De Moor MH, Costa PT, Terracciano A, Krueger RF,De Geus EJ, Toshiko T, Penninx BW, Esko T, Madden PA, Derringer J, Amin N. Meta-analysis of genome-wide association studies for personality. Molecular Psychiatry. 2012 Mar 1;17(3):337–49.

89. Keers R, Coleman JR, Lester KJ, Roberts S, Breen G, Thastum M, Bögels S, Schneider S, Heiervang E, Meiser-Stedman R, Nauta M. A genome-wide test of the differential susceptibility hypothesis reveals a genetic predictor of differential response to psychological treatments for child anxiety disorders. Psychotherapy and Psychosomatics. 2016 Apr 5;85(3):146–58.

